# Predictive metabolomics to decipher plant eco-evolutive tendencies and physiological traits

**DOI:** 10.64898/2026.06.05.730148

**Authors:** Cathleen Mirande-Ney, Santiago Trueba, Adam Rochepeau, Régis Burlett, Pierre Pétriacq, Sylvain Delzon, Yves Gibon, Sylvain Prigent

## Abstract

Plant ecological and evolutionary strategies are shaped by interactions between phylogenetic history and environmental constraints, resulting in leaf and stomatal traits. However, traditional trait-based and phylogenetic approaches often fail to fully explain biochemical mechanisms underlying ecological strategies, particularly for leaf and stomatal traits. Plant metabolomes integrate genetic, physiological, and environmental information and therefore represent a promising intermediate phenotype for investigating links between biochemical diversity, functional traits, and evolutionary patterns.

We analysed metabolomic profiles from 74 plant species with various growth forms and ecological types. Using machine learning approaches, we explored whether metabolic variation could predict plant functional divisions, growth forms and phenological types, but also physiological traits related to drought resistance.

Metabolomic data contained structured information associated with variation in plant functional traits, ecological strategies, and phylogenetic relationships. Machine learning models identified with high accuracy distinct metabolic signatures linked to differences among plant functional divisions, growth forms, phenology, and trait values.

Our study demonstrates that predictive metabolomics provides a powerful and integrative framework to investigate plant ecological and evolutionary strategies. By linking biochemical diversity with plant phylogeny, and ecophysiological traits across multiple species, this approach offers new opportunities to explore the mechanistic basis of plant evolution.

## 1. Introduction

Plants exhibit a wide diversity of ecological strategies, shaped by evolutionary history and adaptation to contrasting environments. These strategies are expressed through coordinated suites of functional traits, including growth form, phenology, and resource-use patterns, which influence plant performance and ecosystem functioning (Violle *et al*., 2007; Díaz *et al*., 2016). Traditional ecological and evolutionary studies have largely relied on phenotypic measurements, trait-based ecology, genetic markers, and phylogenetic comparative methods to infer adaptive strategies (Losos, 2008; Savolainen *et al*., 2013; Zakharova *et al*., 2019). While these approaches provide valuable insights, they often capture only a part of the mechanisms underlying plant functional variation. Physiological traits, particularly leaf and stomatal characteristics, frequently show weak phylogenetic signal due to environmental filtering, phenotypic plasticity, or convergent evolution (Yin *et al*., 2020; Guillemot *et al*., 2022; Ávila-Lovera *et al*., 2023; Trueba *et al*., 2026). Yet, leaf and stomatal traits such as specific leaf area, leaf water content, stomatal area and density, and maximum (*g□□□*) and minimum (*g□□□*) leaf conductance, underpin fundamental carbon–water trade-offs that shape plant life styles along environmental gradients (Wright *et al*., 2004; Franks & Beerling, 2009; Burlett *et al*., 2025).

Metabolomic profiles reflect both lineage-specific biosynthetic pathways and flexible biochemical responses to environmental conditions, integrating downstream consequences of gene expression, physiological performance, and stress responses (Fiehn, 2002; Manickam *et al*., 2023). Therefore, plant metabolism represents a promising integrative phenotype that can capture links between evolutionary history, ecological adaptation, and functional traits. Comparative studies suggest that metabolomic variation can simultaneously reflect phylogenetic relationships and ecological strategies, providing a biochemical framework to link ancestry, adaptation, and functional trait expression (Sardans *et al*., 2011; Montesinos-Navarro *et al*., 2020; Peguero *et al*., 2021; Walker *et al*., 2022, 2023). Despite this potential, the ability of metabolomics to predict physiological traits, plant functional divisions, and phenological strategies across multiple species remains largely unexplored.

Recent advances in high-throughput metabolomics have generated increasingly complex datasets, requiring analytical approaches capable of extracting biologically meaningful patterns (Hajjar *et al*., 2023; Barros Santos *et al*., 2025). Machine learning methods, and more specifically generalised linear models (GLMs), offer powerful tools to identify linear and multivariate relationships within high-dimensional biological data (Dussarrat *et al*., 2022, 2023, 2025). Integrating metabolomics with machine learning thus provides a unique opportunity to test whether biochemical diversity alone can explain variation in key traits and ecological strategies, advancing our understanding of plant adaptation.

Here, we analysed the metabolomes of 74 vascular plant species from 38 families to test whether metabolomic profiles can predict variation in plant functional divisions, growth forms, phenological strategies, and leaf and stomatal traits. This study introduces a multispecies framework that combines metabolomics and machine learning to link biochemical diversity with phylogenetic structure, plant life styles, and leaf and stomatal physiology, thereby advancing our understanding of how metabolic variation underpins plant ecological and evolutionary strategies.

## 2. Materials and methods

### 2.1. Sampling

In total, 74 vascular plant species belonging to 38 plant families (Supporting Information Table S1) were analysed in this study. Angiosperms are particularly well represented with 63 species from major clades of flowering plants. Sampling also covered gymnosperms and ferns with 6 and 5 species, respectively. Sampling was mainly carried out around the campus of the University of Bordeaux (Pessac, France), and supplementary plant material was collected from local nurseries (Table S1). For each species, fully developed leaves of those plants were harvested, five biological replicates corresponding to multiple plants were collected, directly snap frozen in liquid nitrogen, brought back to the laboratory in liquid nitrogen and stored at −80°C until freeze-drying. Sampling was performed from March to April (2023-2024) between 09:30 and 12:30 am. In this study, one epiphyte (*Viscum album*), one vine (*Hedera helix*), and succulent herbs (*Aloe mitriformis* and *Kalanchoe tetraphylla*) were categorised as shrub, tree and herb, respectively, due to insufficient replicates. Althought this classification does not correspond to the growth form of the corresponding species, they correspond to the closest degree of wood production of each species.

### 2.1. Measurements of functional traits, leaf minimum conductance, stomatal features and maximum theoretical conductance

Functional, structural and physiological traits were measured on branches belonging to the same individuals, or the same populations in case of annual herbaceous and small-sized plants, that were used for the metabolomic analyses. Branches were collected and rehydrated overnight using standard protocols (Trueba *et al*., 2019). Rehydrated leaves were collected and immediately weighed with a 4-digit balance (Pioneer, Ohaus, USA) to obtain the water-saturated turgid mass (*M*_t_; in g). Leaves were then scanned (v850 pro, Epson, Japan) to calculate projected leaf area (*A*_leaf_; in m^2^). Sampled leaves were sealed with paraffin to avoid excessive dehydration from the cut ends of the petioles, and tagged. Leaf dehydration was measured using a *droughtbox* device adapted from previously published protocols (Billon *et al*., 2020). Measurements were carried out at stable conditions of 25°C chamber temperature and 60% relative humidity, resulting in a vapour pressure deficit (VPD) of c. 1.27 *k*Pa. Monitoring of weight loss was carried out every 5 minutes to obtain continuous fresh mass (M_f_; in g) values. Measurements ended after reaching important dehydration levels, leaves were then put in an oven at 65°C for 72h and dry mass (*M*_d_; in g) was measured to calculate relative water content during the entire course of dehydration using equation 1:

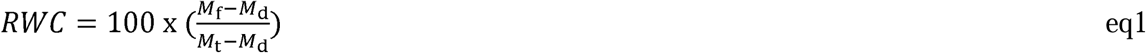

Minimum conductance (*g*_min_; in mmol.m^-2^.s^-1^) was then computed for each sample as the rate of water loss divided by its driving force (VPD) using equation 2:

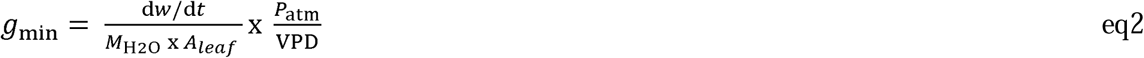

where d*w*/d*t* is the negative slope of the weight∼time relationship from the dehydration curve of the experiment, M_H2O_ is the molecular weight of water (18.01 g mol^-1^), *A*_leaf_ is the projected leaf area (in m^2^), P_atm_ is the atmospheric pressure in the chamber (set to *c*. 101.9 kPa), and VPD is the vapor pressure deficit between the inner leaf and the air inside the chamber (in kPa). We used continuous RWC estimations to measure the rates of water loss between the boundaries of 80% and 50% RWC following recent recommendations (Burlett et al. 2025) to obtain comparable conductance measurements between similar water status thresholds.

Additionally, using the previous measurements, we estimated leaf mass per area as LMA= *M*_d_/*A*_leaf_, and leaf water content as 100 x (M_f_ - M_d_)/ M_d_. We collected stomatal traits on three leaves per species that were adjacent to the leaves sampled for *g*_min_ estimations. We made varnish impressions of both leaf surfaces and image acquisition was done with an optical microscope (Leica DM2500; Leica Microsystems, Buffalo Grove, IL, USA). We estimated stomatal density (*D*, in stomata.m^-2^) and area (*S*, in m^2^). In amphistomatous leaves (i.e. leaves presenting stomatal pores on both sides of the lamina) we averaged both sides to estimate theoretical *g*_max_ (in mmol.m^-2^.s^-1^) based on previously established theories of stomatal conductance based on anatomical traits (Franks & Beerling, 2009) using equation 3:

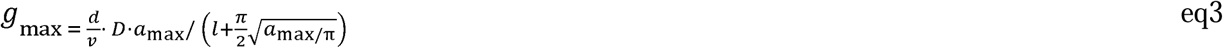

where *d* is the diffusivity of water vapor in air (m^2^·s^-1^), *v* is the molar volume of air (m^3^·mmol^-1^), *a*_max_ and *l* are the maximum area of the open stomatal pore and its depth, respectively, which were set by stomatal area *S* (calculated as guard cell length, *L*, multiplied by width, *W*, of the stomatal pore). All image measurements were performed using *ImageJ* (Schneider *et al*., 2012).

### 2.2. Metabolomics analysis

Freeze-dried plant material (10 mg) was extracted using an ethanol-based solvent system to recover semi-polar metabolites, following Luna *et al*., (2020). Extracts were analysed by untargeted metabolomics using liquid chromatography-high-resolution tandem mass spectrometry (LC-HRMS/MS) on an LTQ-Orbitrap Elite mass spectrometer (ThermoScientific, Bremen, Germany) equipped with an electrospray ionisation (ESI) source operating in negative mode, with data-dependent MS/MS acquisition as described in Dussarrat *et al*., (2022).

Separation was performed on a C18 column (C18-Gemini, 2.0 × 150 mm, 3 µm, 110 Å; Phenomenex, Torrance, CA, USA). MS spectra were acquired over an *m/z* range of 50-1500 with a retention time window of 0-18 min. Data-dependent MS/MS was triggered on the most intense precursor ions using higher-energy collisional dissociation (HCD) at a normalised collision energy of 60 eV. A total of 20 extraction blanks, 66 quality control (QC) samples, and 380 plant samples were injected.

### 2.3. Processing of Metabolomic Datasets

Raw LC-MS data were processed in MS-DIAL v4.9.221218 with optimised parameters: MS1 tolerance 0.008 Da, MS2 tolerance 0.025 Da, minimum peak height 20,000, and retention time tolerance 0.5 min. Peak detection used a linear weighted moving average smoothing algorithm, and features were aligned across samples using QC references (retention time tolerance 0.2 min; mass tolerance 0.015 Da). Background signals were removed using a sample-to-blank ratio ≥ 5, and gap filling was applied to generate a complete data matrix.

Metabolite annotation was performed by matching MS/MS spectra against a curated MSP library (Dablanc *et al*., 2024) and cross-checked with an in-house IROA database, including retention times. Annotated features were assigned chemical ontology using ClassyFire (Djoumbou Feunang *et al*., 2016). Raw signals were normalised to the internal standard methylvanillate (250 µg·mL ¹) and corrected for analytical drift using LOWESS regression based on QC injections every 11 samples. After curation (blank check, signal-to-noise >10, QC CV <30%), 4,725 features were retained for downstream analysis.

Finally, data were normalised by sample weight, log-transformed, and Pareto-scaled using MetaboAnalyst v.6 (Pang *et al*., 2024) prior to multivariate statistical analyses. Extraction blanks, standards, and QC samples were excluded from visualisation for clarity.

### 2.4. Multivariate statistical analyses

Data analysis was performed using R (v4.4.2; R Core Team, 2024) and RStudio (v2024.12.0; RStudio Team, 2024). Data frames manipulation were conducted using R packages “dplyr” (v1.1.4; Wickham et al., 2023), “tibble” (v3.2.1; Müller *et al*., 2023) and “tidyr” (v1.3.1; Wickham et al., 2024b). Feature richness was calculated for each division, growth form and phenology to assess chemical diversity across different phyletic groups and ecological traits (growth form and phenology) from the raw and uncurated metabolomics dataset. Richness was defined as the total number of distinct features detected per species observed within each category. We consider that a feature belongs to a species if it is present in at least 3 replicates. To explore overlap in feature composition among selected plant categories, Venn diagrams were constructed using the “VennDiagram” v1.7.3 package (Chen, 2022) from the uncurated metabolomics dataset. Given the imbalance in the number of species among divisions: 63 angiosperms compared to only 6 gymnosperms and 5 ferns, to construct the Venn diagram for the division category, we generated an average list of angiosperm-specific compounds by randomly sampling 6 angiosperm species 10,000 times. For each iteration, we recorded the detected features and retained only those that appeared in at least 66% of the repetitions, representing consistently present features across subsets of angiosperms. To test for statistically significant differences in feature richness among plant divisions or taxonomic categories, a one-way Kruskal-Wallis test was performed. When significant group effects were detected (*p* < 0.05), post hoc comparisons were conducted using Dunn’s test with Bonferroni correction. Analyses were performed using basic functions in the “rstatix” package v0.7.2 (Kassambara, 2023). Compact letter displays for post hoc group comparisons were generated using the “multcompView” package (v0.1-10; Graves et al., 2024). Figures, including boxplots and barplots, were created using “ggplot2” (v3.5.1; Wickham, 2016; Wickham et al., 2024). Factor levels were reordered and formatted for plotting with “forcats” v1.0.0 (Wickham, 2023). Layout adjustments and multi-plot arrangements were handled using “gridExtra” v2.3 (Auguie and Antonov, 2017). Hierarchical clustering was performed to explore patterns of similarity among plant species based on their average metabolomic profiles. The similarity between species was quantified using Pearson correlation coefficients. Clustering was performed using the Ward’s method (ward.D2) and visualised with the “dendextend” packages v1.19.0 (Galili *et al*., 2024). A phylogenetic tree with species present in the metabolomics dataset was built using the dated mega-tree GBOTB.extended.tre (Smith & Brown, 2018), using the “V.PhyloMaker” package (Jin & Qian, 2019). To assess congruence between clustering based on metabolomic profiles and phylogenetic relationships, tree comparison analyses were performed. Visual comparisons were facilitated using tanglegrams and the “step2side” method, highlighting differences in topology and clustering. Distance-based metrics such as baker correlation coefficient and cophenetic correlation coefficient were calculated using the “dendextend” package v1.19.0; (Galili *et al*., 2024)).

### 2.5. Generalised multilinear models

Ridge, LASSO and Elastic-net models (GLM) were constructed in R with the glmnet package (v 3.0-2; Friedman et al., 2010; Tay et al., 2023). For both classification and regression tasks, GLM was used to model the relationship between metabolomic profiles and, plant taxonomy (genus, species, family, order, division, growth form and phenology) and physiological traits (leaf area, leaf mass per area, leaf water content, stomatal area, stomatal density, maximum theoretical leaf conductance and minimum leaf conductance), respectively. We chose to use GLM, which are sparse linear models, because they are easier to interpret and provide biological insights into the effects of feature importance on taxonomical discrimination. Stratified sampling was performed with a random selection of 75% of the individuals used to construct the models, and by using the remaining 25% to test the quality of the prediction. This process was performed 100 times for each dataset to address the random selection of the training and validation sets. Ten-fold cross-validation was repeated 3 times in the construction of the models to decrease overfitting, and the mean square error was used to select the best models during the training step, based on a grid search over 40×40 combinations of the two penalisation parameters (α and λ), with α values distributed between 0 and 1. For classification tasks, model performance was evaluated using the accuracy. For regression tasks, the quality of the models was assessed on the basis of the coefficient of determination (R²) between measured and predicted values in the test set. Thus, the variables occurring the most (66%) in the models were considered as the most stable predictors of plant taxonomy and physiological traits. The average values for the various physiological traits were consistent across all replicates of a species, as these traits were not measured specifically for our samples.

Furthermore, to assess the generalisability of the GLM models and provide a more realistic evaluation of model performance for both classification and regression, class-based cross-validation was performed using Partial Least Squares (PLS) model. Specifically, a leave-one-class-out cross-validation approach was used, in which all samples from a single class were held out as the test set in each iteration (5), and the model was trained on samples from all remaining classes. For classification, this analysis was performed only with the classes “division”, “growth form”, “phenology”, “order” and “family”. Only order and family classes containing at least six species were included in the analysis. To evaluate the detailed performance of the classification models, confusion matrices were constructed for each iteration of the leave-one-class-out cross-validation. For regression tasks, class-based cross-validation was performed by species using a subset of the most stable predictors of leaf physiological traits (important variables with more than 66% of occurrence in the model, representing 240, 193, 233, 189, 201, 208 and 198 features for leaf area, LMA, water content, stomatal area, stomatal density, *g*_min_ and *g*_max_, respectively), and predicted values for each held-out species were compared to observed values using R². For a final test to check for possible data-leakage, the same approach was used while removing a few species from the dataset at the very beginning of the process and comparing both results on *g*_min_ prediction.

## 3. Results

### 3.1. Phytochemical characterisation of plant species reveals that most metabolites are shared among species

Untargeted metabolomics analysis was performed on 74 plant vascular plant species spanning a wide growth from and phylogenetic diversity, yielding a total of 12,884 metabolic features. To ensure the comprehensive exploration of metabolic diversity, i.e. including signals that might be attenuated or lost in quality control samples due to dilution, uncurated features were analysed without QC-based coefficient of variation filtering for cheminal richness and venn analyses. A great majority of these features, up to 80%, were found to be shared among all species examined. No significant difference was found between angiosperms, gymnosperms and ferns for the number of features (chemical richness, Fig. 1). In contrast, chemical richness was significantly higher in trees compared to herbs and shrubs, with 10% more metabolic features that included compounds of almost all classes. This trend was also noticeable for plant phenology, with woody deciduous and evergreen plants showing greater richness than herbaceous perennials and annuals. Gymnosperms displayed more lipids and lipids-like molecules than angiosperms. Regarding plant phenology, no significant differences were identified between annual and perennial herbs and between woody deciduous and evergreens. However, deciduous and evergreen trees had more features in almost all compound classes compared to annual and perennial herbs.

**Figure 1:**
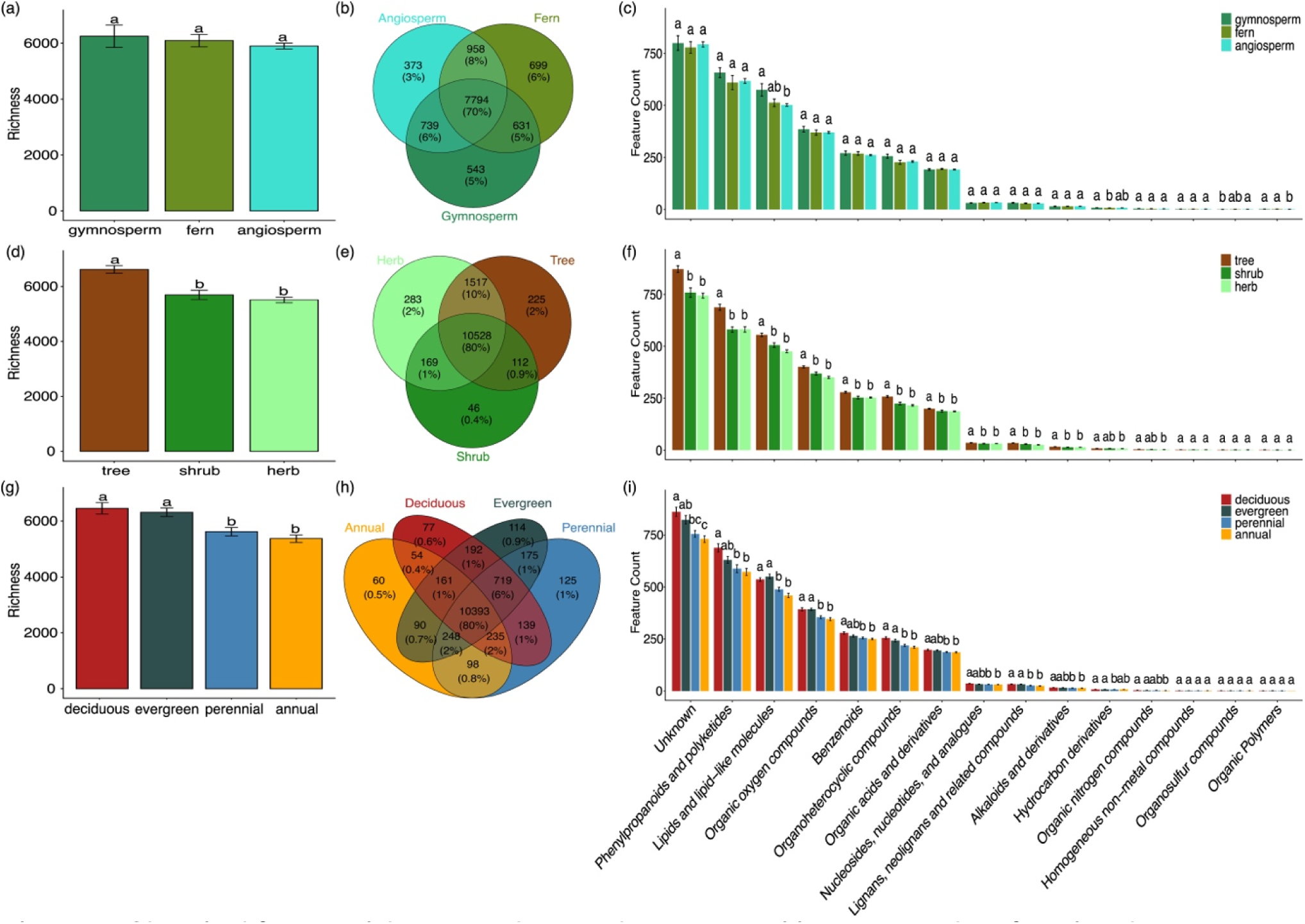
Chemical feature richness and superclass composition across plant functional divisions. (a, d, g) Chemical feature richness (number of detected features) across plant divisions (angiosperms, gymnosperms, ferns), growth forms (herbs, trees, shrubs), and leaf phenology (annual herbs, perennial herbs, woody deciduous, woody evergreen). (b, e, h) Venn diagrams highlighting shared and unique chemical features among categories. (c, f, i) Barplots of average superclass counts per category, revealing differences in chemical class distribution. Statistical differences were tested using Kruskal-Wallis test followed by Dunn’s test with Bonferroni correction; letters indicate significant pairwise differences (α = 0.05).

### 3.2. Metabolomics as a phylogenetics proxy tool

Having established that there are substantial differences in feature occurrence between plant growth forms and phenology classes, we investigated whether metabolomics could be used to predict eco-evolutive trajectories and physiological traits. For this, we used the filtered dataset of 4,725 features to retain the most robust metabolomics signals. Hierarchical clustering allowed species to be sorted relatively well, except for *Ginkgo bilboa* that clustered among angiosperms (Fig. 2). The dendrogram obtained with metabolomics data resembled that of the phylogenetic tree, with an alignment quality of 0.095 and a baker correlation coefficient of 0.21 (Fig. 2).

**Figure 2:**
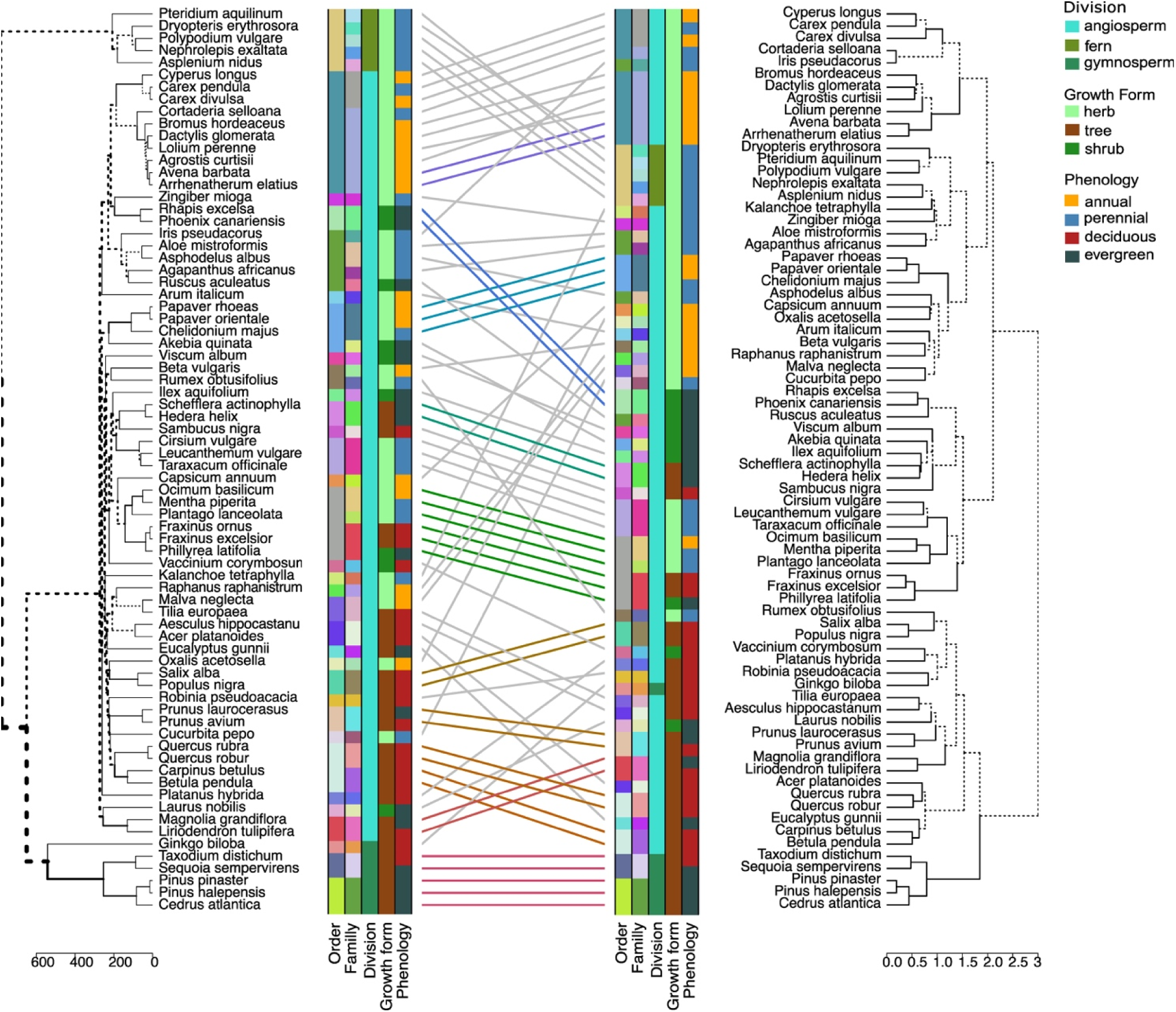
Plant metabolome aligns with phylogeny. Comparison of metabolomic hierarchical clustering (Pearson + Ward.D2) (right) and phylogenetic tree (left). Colored lines between trees link identical species and highlight major phylogenetic groups. Dashed branches indicate clustering uncertainty or lower similarity. The alignment was generated using the dendextend R package with the tanglegram function and step2side method.

Since unsupervised clustering analysis considers all variables, we next deployed supervised, multilinear modelling, which maximises variable discrimination. For each variable to be predicted, 100 models were built with 75% of the observations and then tested with the remaining 25%. Metabolic features used in more than 66 models were considered predictive, and those used in all 100 models were top predictors. The leaf metabolome made it possible to discriminate division, order, family, genus and even species with a high level of accuracy (Fig. S1). Validation of these results was then carried out using PLS model with the most frequently used variables among the 100 models, by building models with all species except one, the latter being then used to verify the accuracy of the model (“leave-one-out”).

Most species were well classified according to phylogeny, except for members of the Asparagales order that were misclassified as Poales, and members of the Cyperaceae family that were misclassified as Poaceae (Fig. S1). The confusion matrix demonstrated a relatively accurate classification, with misclassification rates of 4.1%, 10% and 11.2% for division, order and family, respectively. The use of GLMs also made it possible to distinguish gymnosperms from angiosperms among trees, with an accuracy of 0.99 and 160 features (Fig. S2; Supplementary Table 2). Among the annotated best predictors were phenylalanine, shikimate, metabolites of dicarboxylic metabolism and diterpenoid synthesis, which were positively associated with gymnosperms. Besides, isoflavonoids, succinic acid, N-acetylphenylalanine and coumarins were positively associated with angiosperms. Similarly, GLM prediction was able to distinguish ferns from angiosperm herbs with an accuracy of 1 and 107 features (Fig. S2; Supplementary table 3). Positive predictors of ferns included prenol lipids, arachidonic acid, jasmonic acid, quinic acid, shikimic acid and flavonoids, whereas positive predictors of angiosperm herbs included raffinose and serine.

### 3.3. Metabolome-based prediction of plant growth forms

Multilinear modelling made it possible to distinguish trees, shrubs and herbs relatively well, with the confusion matrix showing a misclassification rate of only 11.8%. We identified 209, 132 and 195 metabolic features making an important contribution to the prediction for herbs, shrubs and trees, respectively (Fig. 3; Supplementary Tables 4-6). For herbs, top positive predictors included fumarate, a fatty acid and phosphoryl-choline, while top negative predictors included 4 lipid (lipd-like) compounds, UTP, 2 carbohydrate conjugates and, a depside or depsidone compound. For shrubs, top positive predictors included 3 lipid (lipid-like) compounds, a nucleoside, an organic acid and a carbohydrate conjugate, while top negative predictors included two carboxylic acids and derivatives and xylitol. For trees, most predictors were contributing positively to the model. With the exception of an amino acid, all top predictors were lipid (lipid-like), benzenoids and secondary metabolites (quinic acid, phenylpropanoids, lignans). Overall, it seems that while herbs were rather characterised by primary metabolites and trees by secondary ones, shrubs appeared in between. Interestingly, luteolin-7-O-glucoside had a negative effect and pinoresinol and quinic acid a positive effect on tree species prediction, whereas they had opposite effects on herb species prediction. Similarly, 2-methoxy cinnamic acid had a negative effect on shrub prediction but a positive effect on tree prediction, and traumatic and fumaric acids had a negative effect on shrub prediction but a positive effect on herb prediction.

**Figure 3:**
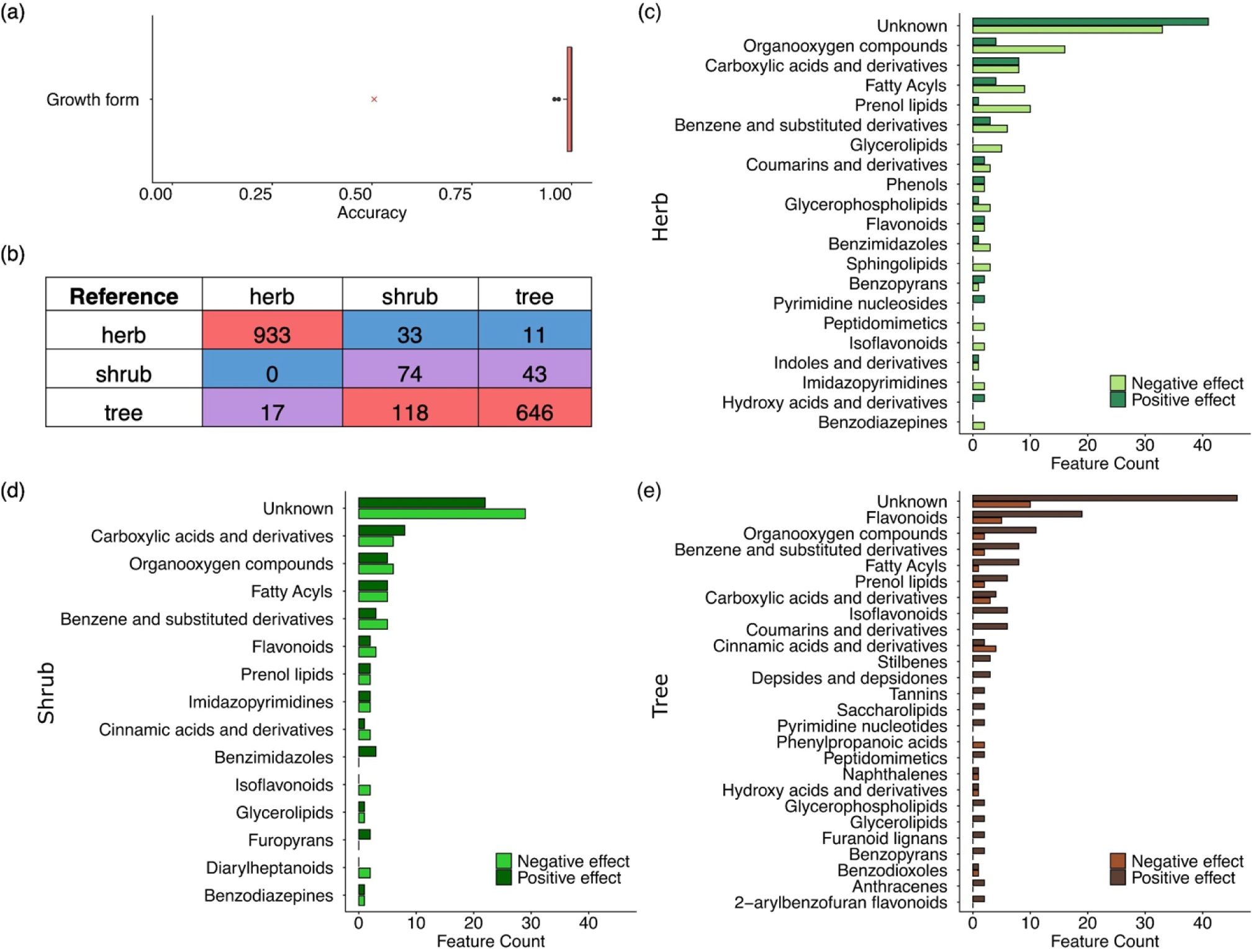
(a) Accuracies obtained by predicting plant growth form using GLM models based on leaf metabolome. Red crosses correspond to accuracies obtained with random data. (b) Confusion matrices obtained by predicting plant growth form using PLS models with a leave-one-class-out cross-validation approach based on leaf metabolome. Main compound classes identified by GLM models (≥66% of occurence) for growth form prediction, including (c) herbs, (d) shrubs, and (e) trees. Bar colors represent the effects of the variables on classification (positive or negative) based on the sign of GLM coefficients.

### 3.4. Metabolome-based prediction of phenological classes

For consistency, the analysis was only conducted with angiosperms (there was poor sampling and only one phenological class for ferns or gymnosperms). Epiphytes, vines and succulent herbs were also excluded due to insufficient replication.

GLM prediction distinguished evergreen from deciduous trees with an accuracy of 0.99, with 86 important features (Fig. 4; Supplementary Table 7). The top positive predictor of the evergreen character was a sesquiterpenoid, whereas for deciduous tree it was a benzophenone. The proportion of lipids (lipid like molecules) and organic oxygen compounds (including carbohydrates, alcohols, amino acids, and respective derivatives) was higher among predictors of the evergreen character, whereas the proportion of benzenoids and phenylpropanoids was higher among predictors of the deciduous character.

**Figure 4:**
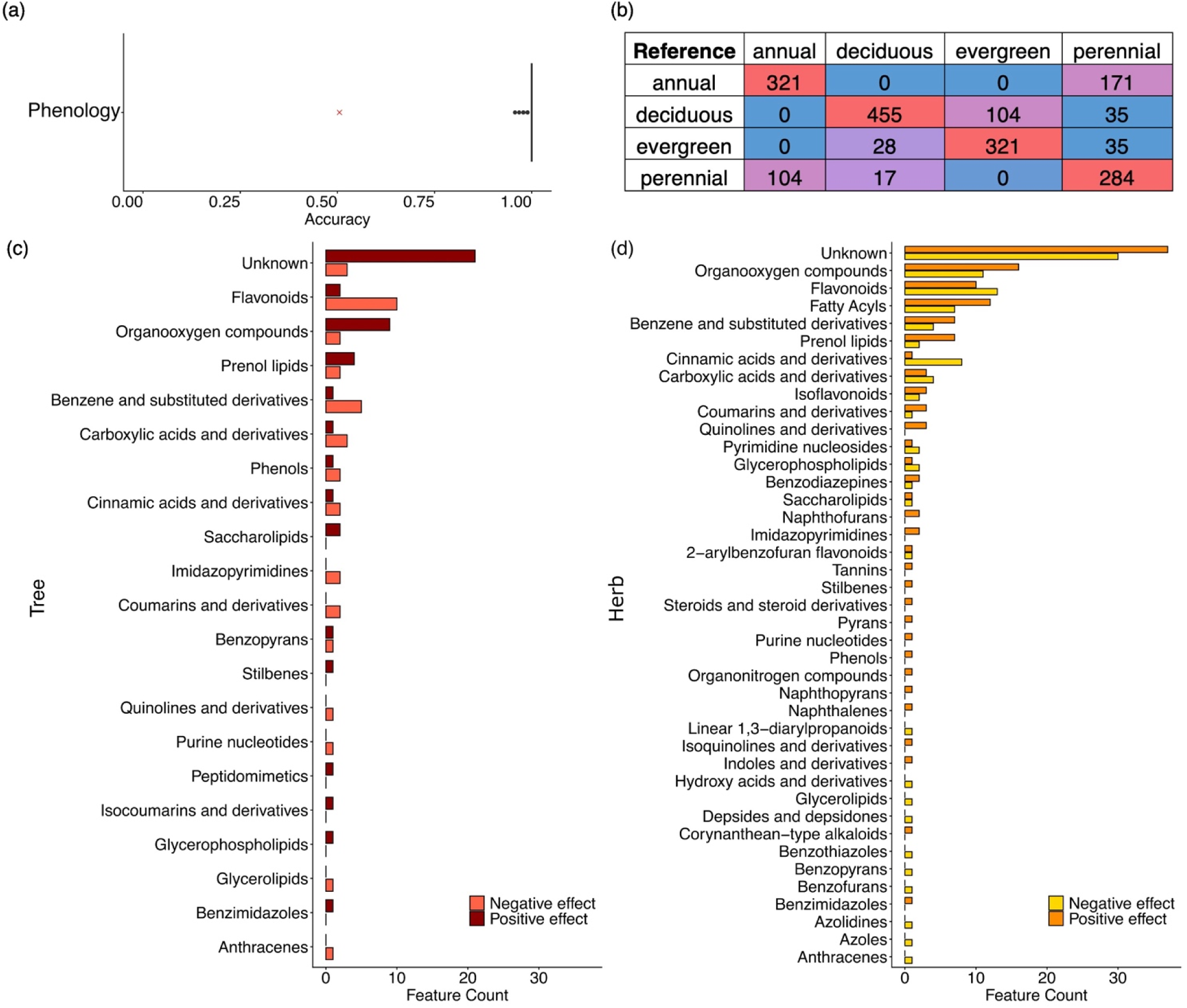
(a) Accuracies obtained by predicting plant phenology using GLM models based on leaf metabolome. Red crosses correspond to accuracies obtained with random data. (b) Confusion matrices obtained by predicting plant phenology family using PLS models with a leave-one-class-out cross-validation approach based on leaf metabolome. Main compound classes identified by GLM models (≥66% of occurence) allowing (c) woody evergreen classification against woody deciduous and (d) perennial herbs classification against annual herbs. Bar colors represent the effects of the variables on classification (positive or negative) based on the sign of GLM coefficients.

GLM prediction was also able to distinguish between annual and perennial herbs with an accuracy of 0.99 and with 224 predictors (Fig. 4; Supplementary Table 8). Top positive predictors of perennial herbs included terpene glycosides and conjugates of carbohydrates. It is also worth mentioning the polyols maltitol and myoinositol, the flavonoids phloridzin, bavachinin and acacetin, and the stilbene resveratrol among the 125 positive predictors of the perennial character. Top negative predictors were a cyclic ketone and the flavonol isorhamnetin. Glutamine, glutathione and fucose-1-phosphate were also found among the 99 negative predictors. More globally, more lipids (lipid-like molecules) and organoheterocyclic compounds were predictors for the perennial character and more phenylpropanoids and polyketides for the annual character.

### 3.5. Metabolome-based prediction of leaf structural traits

Evergreen trees were characterised by low leaf area, high leaf mass per area (LMA) and low water content, whereas deciduous trees showed low LMA, high leaf area and high water content. Herbs (annual and perennial combined) also showed low LMA and high water content, but a low to moderate leaf area (Fig. 5). Predictions of leaf area, LMA and water content were highly accurate with R^2^ values reaching 0.96 for these traits when the leave-one-out approach was used (Fig. 5 and Fig. S3-S4). For leaf area, top positive predictors (out of 129) included adenosine, benzenoids, and phenylpropanoids such as fraxetin and isobavachin, and top negative ones (out of 111) benzenoids and isoflavonoids (Fig. S3; Supplementary Table 9). For LMA, among the 121 positive predictors, top ones included a benzenoid, a long fatty acyl alcohol, C20- and C19 gibberellins and shikimic acid, while top negative predictors (out of 72) included flavonoids, isoflavonoids and fatty acyls (Fig. 5; Supplementary Table 10). The lipid (lipid-like) superclass was clearly more represented among the negative predictors (14%) than among the positive ones (5%). For water content, we found 99 positive predictors and 134 negative ones (Fig. S4; Supplementary Table 11). Top positive ones included fumaric acid and top negative ones, an amino acid. There were more than 2-times more negative predictors than positive ones among lipids (lipid-like) and organic oxygen compound classes. Interestingly, only trehalose, two prenolic lipids and one isoflavonone were common predictors for all three traits.

**Figure 5:**
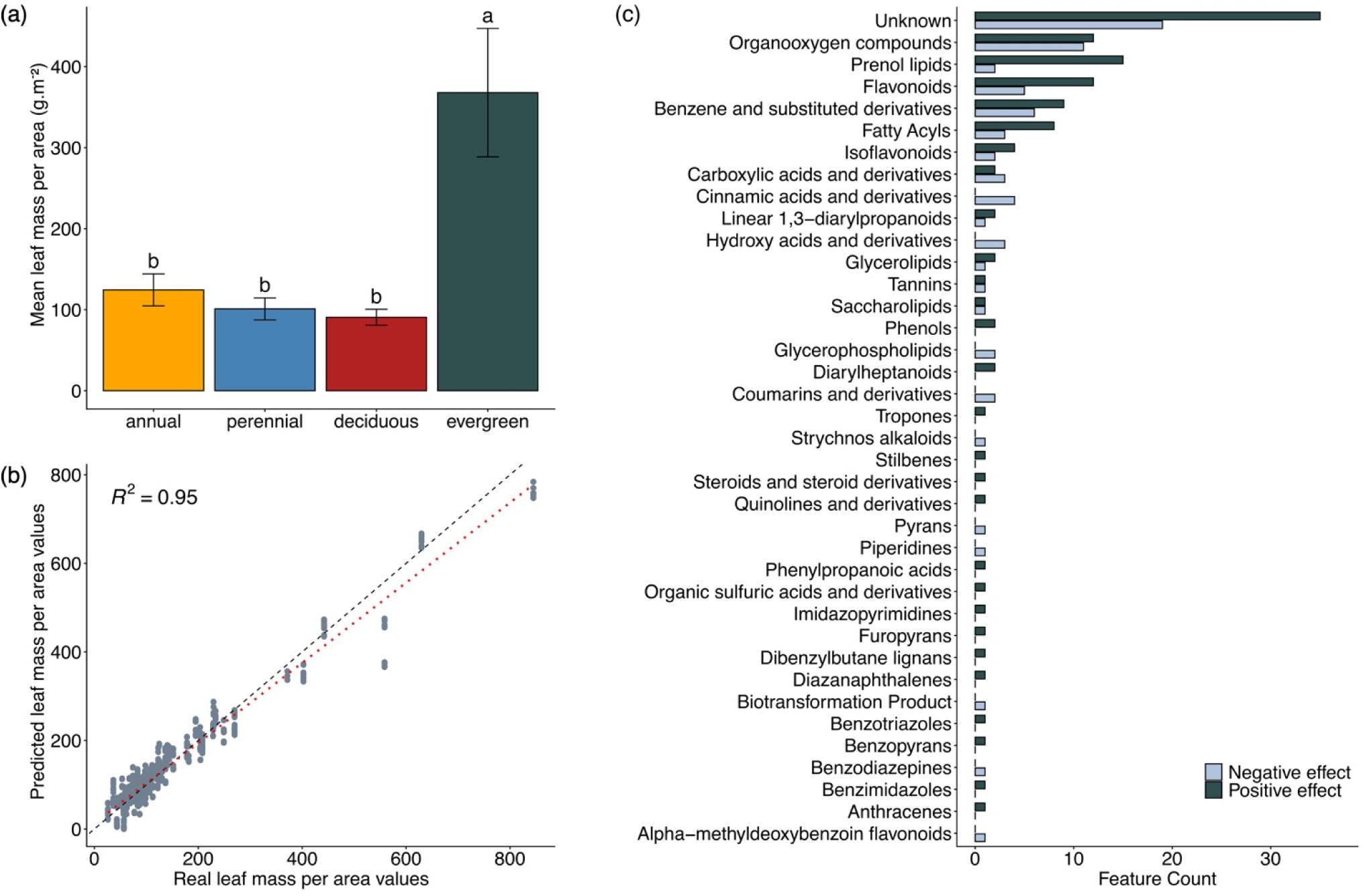
a) Leaf mass per area (LMA) variation across plant leaf phenological types (annual and perennial herbs, and deciduous and evergreen trees). LMA is expressed in grammes by squared metres (g.m^-^²). Statistical differences were assessed using a Kruskal–Wallis test followed by Dunn’s post hoc test with Bonferroni correction; different letters indicate significant pairwise differences (α = 0.05). (b) Predictive metabolomic performance for leaf area obtained using PLS models with a species-based cross-validation approach, based on a subset of the most stable predictors identified by GLM (i.e., variables occurring in more than 66% of models). The red dashed line represents the linear fit between real and predicted values, while the black dashed line indicates the 1:1 relationship; the coefficient of determination (R²) reflects the goodness-of-fit of this relationship. (c) Main compound classes identified by GLM models (occurrence ≥ 66%) contributing to LMA prediction. Bar colors represent the effects of the variables on regression (positive or negative) based on the sign of GLM coefficients.

### 3.6. Metabolome-based prediction of leaf stomatal and hydraulic traits

Stomatal area, stomatal density, *g*_min_ and *g*_max_ varied dramatically between species (Fig. 6). Thus, evergreen trees showed high stomatal area and *g*_max_ but relatively low stomatal density and *g*_min_, whereas deciduous trees showed low stomatal area but higher stomatal density and *g*_max_. Herbs showed moderately low stomatal area, density and *g*_max_ but high *g*_min_.

**Figure 6:**
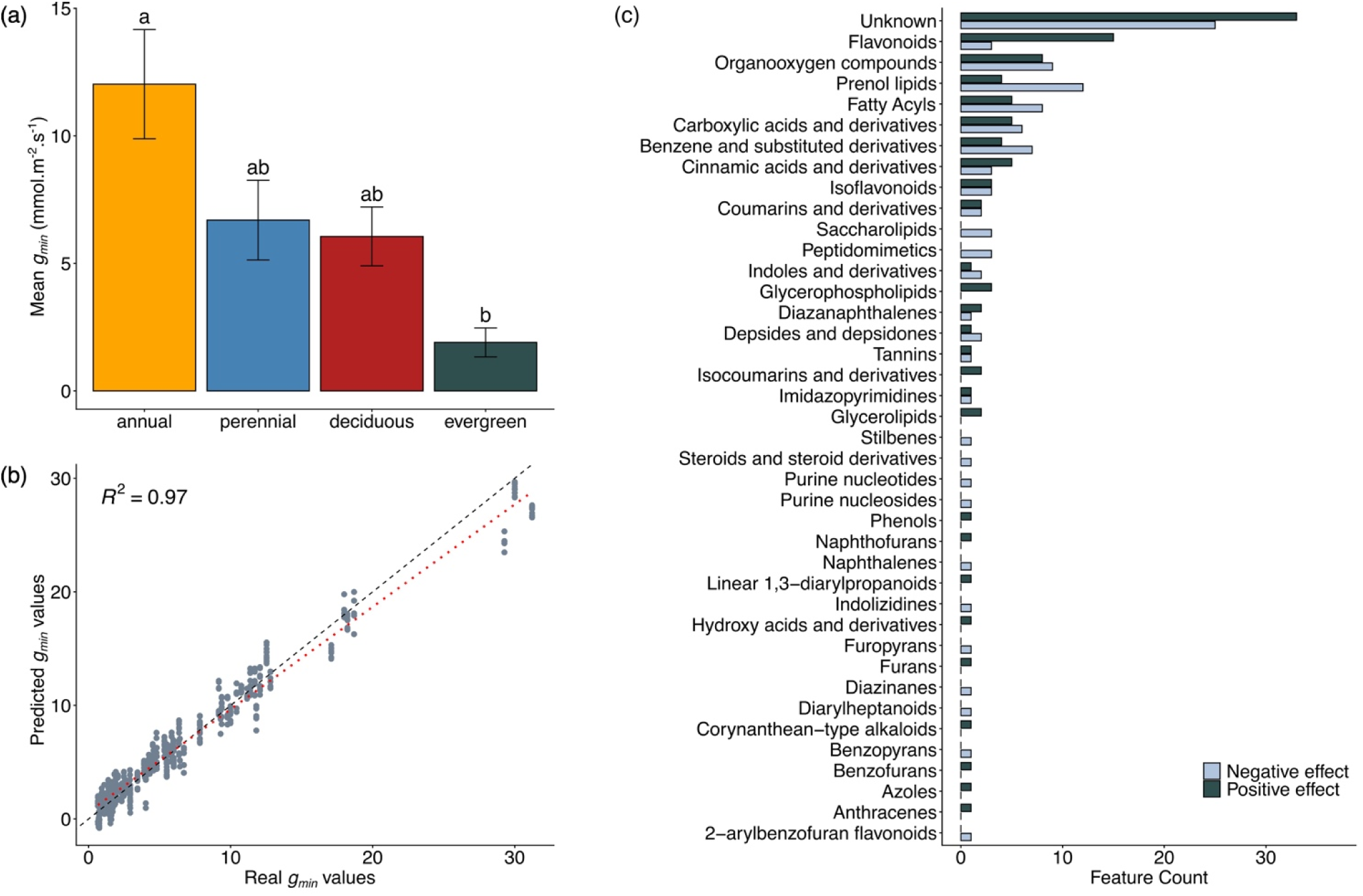
Minimum conductance (*g*_min_) variation across plant leaf phenological types (annual and perennial herbs, and deciduous and evergreen trees). *g*_min_ is expressed in millimoles by squared metres by seconds (mmol.m^-^².s^-1^). Statistical differences were assessed using a Kruskal–Wallis test followed by Dunn’s post hoc test with Bonferroni correction; different letters indicate significant pairwise differences (α = 0.05). (b) Predictive metabolomic performance for *g*_min_ obtained using PLS models with a species-based cross-validation approach, based on a subset of the most stable predictors identified by GLM (i.e., variables occurring in more than 66% of models). The red dashed line represents the linear fit between real and predicted values, while the black dashed line indicates the 1:1 relationship; the coefficient of determination (R²) reflects the goodness-of-fit of this relationship. (c) Main compound classes identified by GLM models (occurrence ≥ 66%) contributing to *g*_min_ prediction. Bar colors represent the effects of the variables on regression (positive or negative) based on the sign of GLM coefficients.

Predictions of stomatal area, stomatal density, *g*_min_ and *g*_max_ were very accurate with R^2^ values above 0.96 when the leave-one-out approach was used (Fig. 6 and Fig. S5-S7), with respectively 189, 201, 208 and 198 key contributing features (Tables S12-S15). On the independant final validation set with species removed from the very beginning, average relative error was 2.5 times higher than those observed on the original dataset, with errors reaching a maximum of 8%. For stomatal area, top positive predictors included benzenoids, organooxygen compounds, uracil and phenylpropanoids while top negative predictors terpenes, derivatives of organic acids and conjugates of carbohydrates (Fig. S5; Supplementary Table 12). For stomatal density, top positive predictors included derivatives of benzoic acid, glycerophospholipids, phosphate esters, conjugates of carbohydrates and phenylpropanoids, and top negative predictors chlorobenzoic acid and gibberellic acid (Fig. S6; Supplementary Table 13). For *g*_max_, top positive predictors included benzoic acid derivatives, lineolic acid, suberic acid, carbohydrate conjugates and phenylpropanoids, and top negative predictors were gibberellic acid and the phenylpropanoids dipteryxin (Fig. S7; Supplementary Table 14). Finally, for *g*_min_, top positive predictors included glycerophospholipids, derivatives of carboxylic acids, conjugates of carbohydrates and phenylpropanoids, and top negative predictors included a terpene glycoside, isocitrate and further phenylpropanoids (Fig. 6; Supplementary Table 15). More globally, elevated *g*_min_ values appeared to be primarily influenced by the accumulation of glycerophospholipids and phenylpropanoids, such as flavonoids, cinnamic acids, or isocoumarins, while lower *g*_min_ values were predominantly associated with the accumulation of lipids, including saccharolipids, prenol lipids, or fatty acyls.

## 4. Discussion

Metabolomics provides an ecologically meaningful dimension in addition to traditional plant phylogenetics, which has largely relied on DNA sequences and morphological traits. By analysing leaves from 74 vascular plant species using LCMS and bioinformatics tools, we found that metabolic profiles of those species were primarily structured by phylogenetic groupings (division, order, family, genus), but also reflected ecological strategies like growth form and phenology (Lee *et al*., 2020; Schweiger *et al*., 2021). Despite 80% of metabolites being shared across species, suggesting possible convergent evolution (Pichersky & Lewinsohn, 2011), specific compounds (<6%) were associated with particular lineages, growth forms or phenology (Fig. 1) as already seen in previous studies (Boachon *et al*., 2018; Schweiger *et al*., 2021).

### Leaf metabolomes reflect plant taxonomy, forms and strategies

Gymnosperms were characterised by high levels of diterpenoids, shikimate derivatives, and arachidonic acid, that are markers of conserved resin-based defences and fatty acid signalling (Wolff *et al*., 2000; Savchenko *et al*., 2010; Hall *et al*., 2013). Angiosperms, by contrast, showed a richer diversity in phenylpropanoids and organic acids involved in growth and metabolic flexibility (Klein & Ramon, 2019; Yao *et al*., 2021; Miyazawa *et al*., 2023, 2025; Piechowiak & Balawejder, 2025). Notably, although phylogenomics places *Ginkgo biloba* firmly within gymnosperms (Wu *et al*., 2013; Yang *et al*., 2024), its metabolomic profile clustered more closely with angiosperms, suggesting convergent chemical strategies possibly linked to shared traits such as leaf and fleshy fruit-like structures, and aspects of sporangial development.

Ferns and herbaceous angiosperms also revealed distinct metabolic strategies. Ferns exhibited high levels of conserved metabolites such as shikimic, quinic, and 3-coumaric acids (de Vries *et al*., 2021; Maeda & Fernie, 2021), alongside arachidonic acid and 5(S)-HETE, that are rarely found in angiosperms but reflect ancestral oxylipin signalling (Jamieson & Reid, 1975; Schluttenhofer, 2020). Ferns also showed unique signatures of microbial or exogenous compounds, like aspterric acid and monoethyl phthalate, potentially due to gene transfer or symbiotic associations (Sun *et al*., 2015; Li *et al*., 2018). Interestingly, ferns shared more metabolic features with angiosperms (8%) than with gymnosperms (5%), despite earlier divergence time. This could possibly reflect adaptations of ferns to niches dominated by angiosperms (Schneider *et al*., 2004). In contrast, angiosperm herbs displayed fewer predictive features but included raffinose and L-serine, consistent with distinct carbon allocation and nitrogen assimilation strategies (McCaskill & Turgeon, 2007).

Trees differ from herbs and shrubs in their chemical composition, with a greater diversity of metabolite classes among their markers, including lignans and phenylpropanoid-derived compounds like flavonoids, coumarins, and cinnamic acids (Hathway, 1962). Trees also showed high levels of carbohydrates and amino acids, consistent with their slower growth rates (Myers & Kitajima, 2007; Poorter & Kitajima, 2007; Atkinson *et al*., 2012). These patterns suggest a conservative metabolic strategy centred on growth-defence trade-offs. Among angiosperm trees, woody evergreen species were the ones that most resembled gymnosperms, as both groups showed high levels of glycosyl conjugates and terpenes, which are markers of structural and stress-related metabolism, while woody deciduous species showed higher levels of specialised metabolites like flavonoids and coumarins, probably reflecting a tendency toward fast defence strategies (Givnish, 2002; Keel & Schädel, 2010).

In contrast, herbaceous plants exhibited metabolic profiles in line with rapid growth and development, especially in annual species. Those species were indeed characterised by high levels of TCA intermediates, nitrogen-rich amino acids like threonine, and pyrimidine nucleosides, hence suggesting fast energy production and biosynthesis (Moffatt & Ashihara, 2002; Zhang & Fernie, 2018). Annual herbs were also richer in glycosylated compounds and cinnamic acid derivatives, which are involved in structural flexibility and defence (Garnier, 1992; Schilmiller *et al*., 2009; Steenackers *et al*., 2019), as well as in sulfated galactolipids and glycerophospholipids that enhance membrane remodelling and stress signalling (Welti *et al*., 2002; Cowan, 2009; Bolik *et al*., 2022). Perennial herbs, on the other hand, were enriched in metabolites involved in long-term resilience, such as osmoprotectants like maltitol and myo-inositol (Moing, 2000; Valluru & Van den Ende, 2011), terpenes, benzenoids, and oxidised fatty acids that support structural integrity and defence (Pichersky & Gershenzon, 2002; Pollard *et al*., 2008).

### Leaf metabolomes enable the prediction of functional traits

Our study shows that plant functional traits, including leaf mass per area (LMA), leaf area, and leaf water content, are deeply embedded in plant metabolism, with metabolic profiles reflecting both growth strategies and structural or defence investments (Gago *et al*., 2016; Schweiger *et al*., 2021; Walker *et al*., 2022, 2023). Fast-growing, low-LMA species appeared enriched in primary metabolites such as amino acids, organic acids, sugars, and nucleotides, which suggests high metabolic activity, energy production, and protein biosynthesis. In contrast, high-LMA species, such as evergreen trees, were found to accumulate structural and defence-related metabolites, including lignin, phenolics, terpenoids, and lipids, which contribute to tissue density, antioxidative protection, and stress resilience (Wright *et al*., 2004; Poorter *et al*., 2009).

Beyond classical leaf traits, our analyses highlight the predictive power of metabolomics for stomatal traits and plant water use. Thus, flavonoids, benzenoids, prenol lipids, and sorbitol-6-phosphate were associated with stomatal traits (Table S12-S13), the latter likely exerting an indirect effect under drought conditions (Fraser *et al*., 2009; Zhao *et al*., 2014; Zhou *et al*., 2017). Leaf metabolite profiles, including sugars, organic acids, and intermediates such as myo-inositol and shikimate, correlate with stomatal conductance, mesophyll conductance and photosynthetic capacity (Daloso *et al*., 2016; Gago *et al*., 2016).

Importantly, we obtained very good predictions for maximum (*g*_max_) and minimum (*g*_min_) leaf conductance, two key traits for plant water use and drought responses. Thus, *g*_max_, the theoretical upper limit of stomatal water vapour diffusion, was associated with metabolites involved in structural development (lignin precursors, lignans, terpenoids) and stomatal regulation (phosphoric esters, glucoraphanin, N-acetyl-4-aminobutyrate, 4-hydroxy-benzoate) (Barkosky & Einhellig, 2003; Lee & and Lee, 2008; Rui & Anderson, 2016; Shtein *et al*., 2017; Pichersky & Raguso, 2018; Zörb *et al*., 2022; Salehin, 2024). High *g*_max_ species, especially deciduous trees, also accumulate phenylpropanoids, supporting ROS modulation and ABA-mediated regulation (Fig. 6c) (Watkins *et al*., 2014, 2017; Kalariya *et al*., 2021). Conversely, *g*_min_, which is lower in woody evergreen species (Trueba *et al*., 2026) and represents residual water losses after stomatal closure, was negatively correlated with lipid-related metabolites that enhance cuticular wax formation and leaf hydrophobicity (Fig. 6 (d)) (Riederer & Schreiber, 2001; Duursma *et al*., 2019; Burlett *et al*., 2025). This is in line with the fact that *g*_min_ negatively correlates with cuticle thickness (Liao *et al*., 2025). Conversely, *g*_min_ was positively correlated with phenylpropanoids and glycerophospholipids, which are involved in stress responses rather than water conservation, suggesting a trade-off between water conductance and defence against abiotic and biotic stress. The *g*_min_ trait plays a critical role in plant survival to extreme droughts (Duursma *et al*., 2019). Its assessment has made significant progress (Burlett *et al*., 2025) but remains relatively tedious. The ability to predict the value of this trait from the leaf metabolome and models built with subsets of species or genotypes, and the direct method could significantly increase the throughput of screens aimed at finding resistant species or varieties. This could be particularly useful for forestry, which faces a dramatic increase in tree mortality due to climate change (Allen *et al*., 2010). More generally, the possibility to easily predict values for various traits, such as those tested here, in particular so-called “hard” functional traits, which evaluation is work-intensive (Belluau & Shipley, 2018), could be extended to a range of further traits.

## 5. Conclusion

The fact that a large proportion of metabolites are shared among a large diversity of species and that a number of predictions could be obtained with them opens a range of new perspectives. Thus, the possibility to predict important traits such as ecophysiological traits related to water use and drought stress resistance could prove very useful in breeding programs aiming at releasing varieties resistant to extreme drought. Also, predictions could be established for many more traits of interest using species diversity, the advantage being that high diversity translates into high dynamics regarding the range of trait values and metabolite concentrations. Indeed, high dynamics imply that the number of observations does not need to be very large. Accordingly, excellent predictions were obtained here, with a relatively small number of observations. As mentioned above, not all annotations are reliable due to the still-limited number of spectral libraries available for plant-derived compounds. While the total number of metabolites produced by the plant kingdom is estimated to be between 50,000 and 100,000 (Alseekh & Fernie, 2023), only a few thousand metabolites seem to be common to all species, including approximately 1,000 primary metabolites. Prioritising the annotation of this core group of common metabolites could be worthwhile. Indeed, high-quality annotation combined with a large number of observations (species diversity and range of environmental conditions) would ultimately allow us to elucidate the functions of these metabolites using top-down modelling approaches.

## 6. Acknowledgements

We thank Pepinière Lelann for graciously providing us with ornamental plant samples. We thank colleagues Malo Leboulch, Cédric Cassan, Georges Randriafanomezantsoa-Radohery, Millena Barros-Santos and Camille Ziegler for their help during field sampling. We thank Jennifer Dudit for assistance in graminoid species identification. We thank Fabienne Soulet for providing us with liquid nitrogen. Metabolomic analyses were enabled through the MetaboHUB infrastructure (MetaboHUB ANR-11-INBS-0010; MetEx+ ANR-21-ESRE-0035; MetaboHUB (JVCE) ANR-24-INBS-0012), and by the PHENOME infrastructure (ANR-11-INBS-0012), funded by the Agence Nationale de la Recherche under the France 2030 program. This study received financial support from the French government in the framework of the IdEX Bordeaux University “Investments for the Future” program/GPR Bordeaux Plant Sciences.

## 8. Credit authorship contribution statement

CMN, SP and YG designed the study; RB, SD and ST designed protocols and sampling for ecophysiology and functional trait measurements. CMN, ST and AR carried out experiments and processed the samples; CMN performed data analyses; CMN, SP and YG wrote the first version of the manuscript; all authors contributed to manuscript revisions.

## 9. Data availability

Metabolomic data are available from the MetaboLights database under the accession number REQ20250521210652.

