## Supplementary material for "Predictive metabolomics to decipher plant eco-evolutive tendencies and physiological traits": Fig.S1

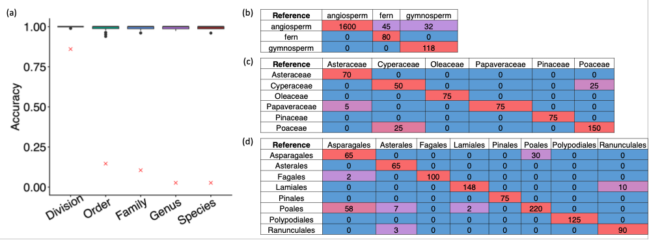

Figure S1: Accuracies obtained by predicting (a) plant taxonomy (division, order, family, genus and species) using GLM models based on leaf metabolome. Red crosses correspond to accuracies obtained with random data. Confusion matrices obtained by predicting plant species (b) division, (c) order, (d) family using GLM models with a leave-one-class-out cross-validation approach based on leaf metabolome.
