## Supplementary material for "Predictive metabolomics to decipher plant eco-evolutive tendencies and physiological traits": Fig.S2

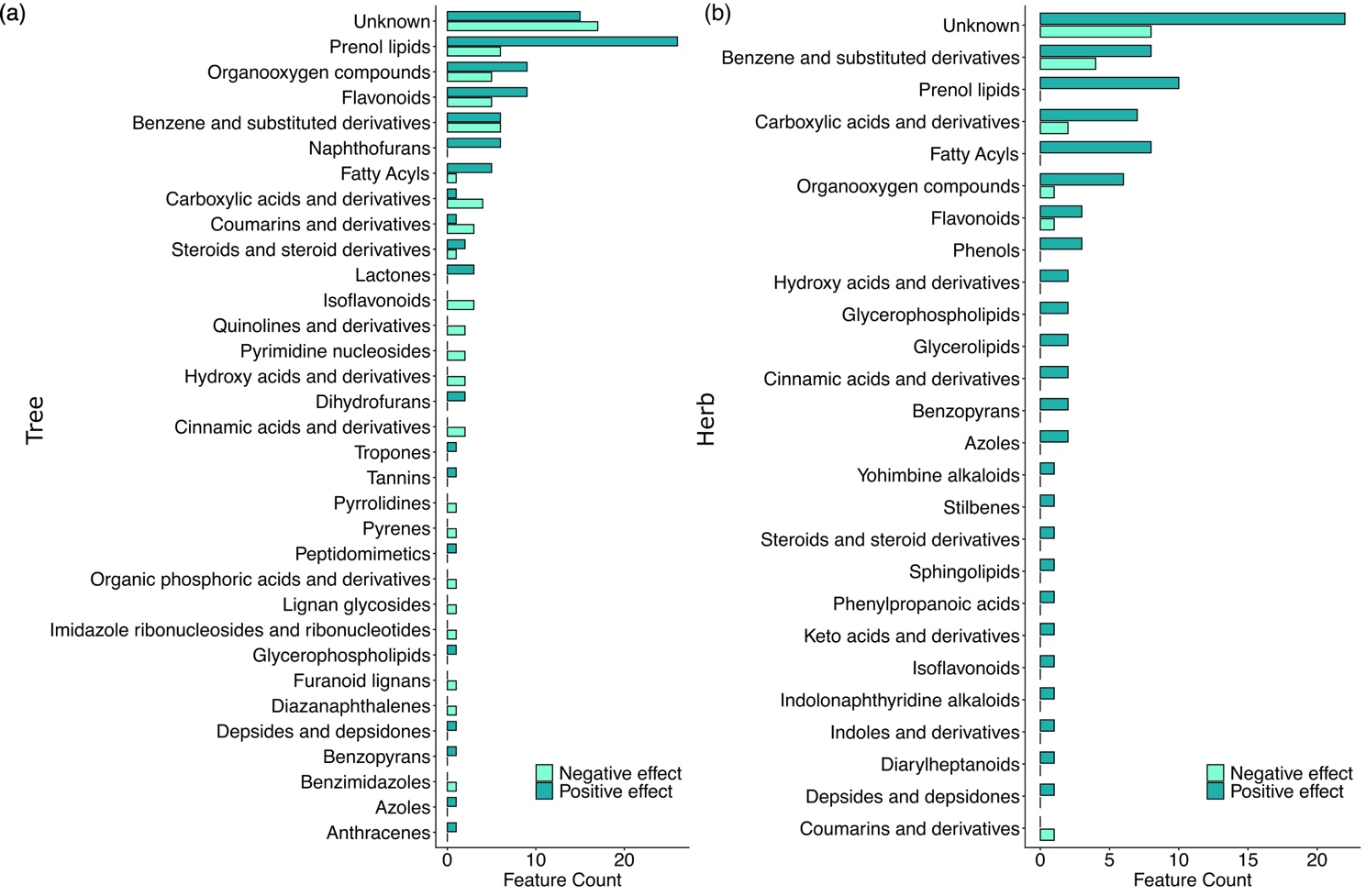

Figure S2: Main compound classes identified by GLM models (≥66% of occurence) allowing (a) gymnosperm tree classification against angiosperm and (b) herbaceous fern classification against angiosperm. Bar colors represent the effects of the variables on classification (positive or negative) based on the sign of GLM coefficients.
