## Supplementary material for "Predictive metabolomics to decipher plant eco-evolutive tendencies and physiological traits": Fig.S7

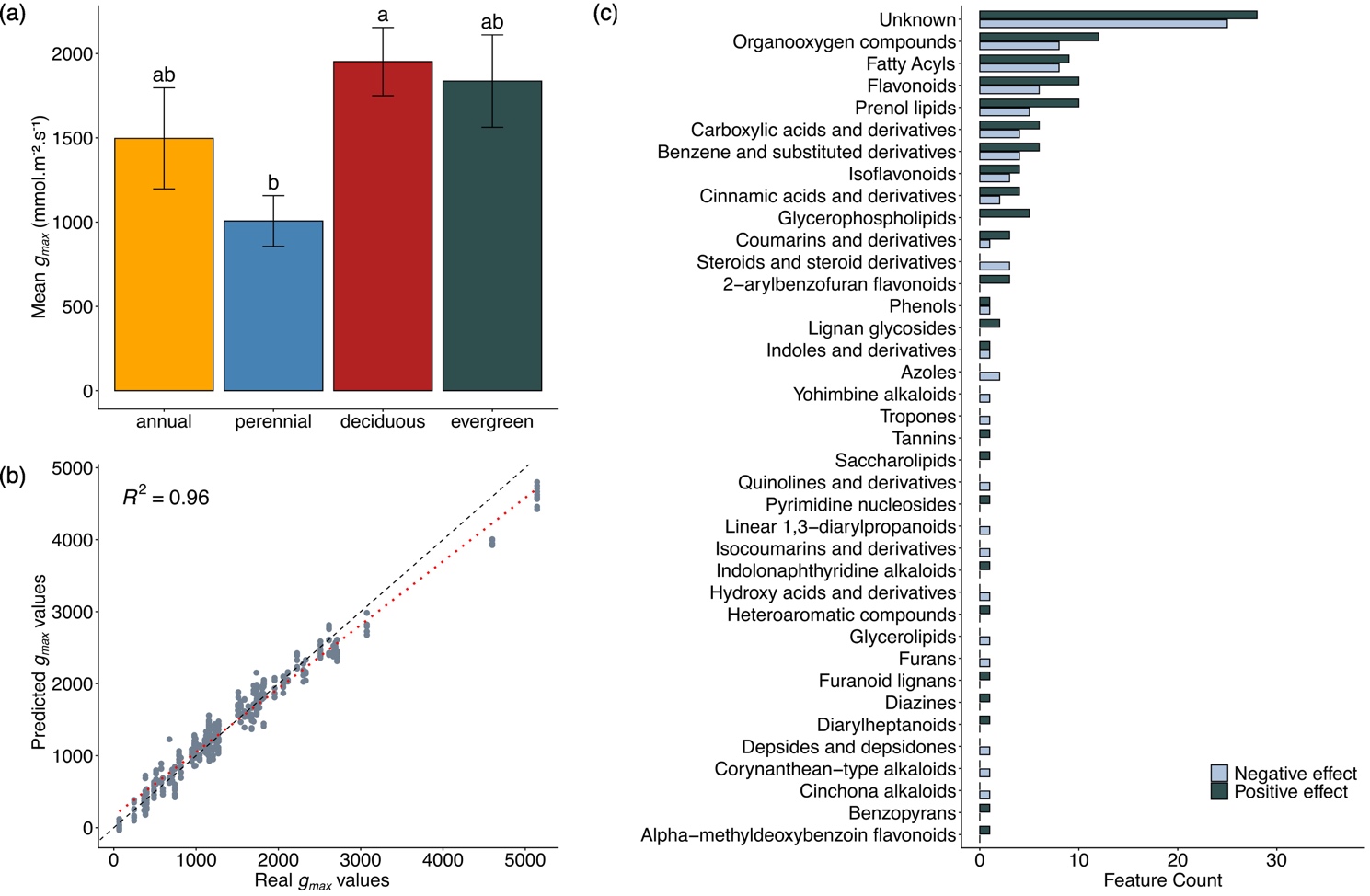

Figure S7: a) Maximum conductance (*g*_max_) variation across plant leaf phenological types (annual and perennial herbs, and deciduous and evergreen trees). *g*_max_ is expressed in millimoles by squared metres by seconds (mmol.m^-^².s^-1^). Statistical differences were assessed using a Kruskal–Wallis test followed by Dunn’s post hoc test with Bonferroni correction; different letters indicate significant pairwise differences (α = 0.05). (b) Predictive metabolomic performance for *g*_max_ obtained using PLS models with a species-based cross-validation approach, based on a subset of the most stable predictors identified by GLM (i.e., variables occurring in more than 66% of models). The red dashed line represents the linear fit between real and predicted values, while the black dashed line indicates the 1:1 relationship; the coefficient of determination (R²) reflects the goodness-of-fit of this relationship. (c) Main compound classes identified by GLM models (occurrence ≥ 66%) contributing to *g*_max_ prediction. Bar colors represent the effects of the variables on regression (positive or negative) based on the sign of GLM coefficients.
